# Nitric oxide radicals are emitted by wasp eggs to kill mold fungi

**DOI:** 10.1101/495085

**Authors:** Erhard Strohm, Gudrun Herzner, Joachim Ruther, Martin Kaltenpoth, Tobias Engl

## Abstract

Detrimental microbes caused the evolution of a great diversity of antimicrobial defenses in plants and animals. Here we show that the eggs of a solitary digger wasp, the European beewolf *Philanthus triangulum,* emit large amounts of the gaseous free radical nitric oxide (NO˙) to protect themselves and their provisions, paralyzed honeybees, against mold fungi. Despite the extraordinary concentrations of nitrogen radicals (NO˙ and its oxidation product NO_2_˙) in the brood cells (~1500ppm), NO˙ is synthesized from L-arginine by an NO-synthase (NOS) as in other animals. The beewolf *NOS* gene revealed no conspicuous differences to related species. However, due to alternative splicing, the NOS-mRNA in beewolf eggs lacks a 144bp exon near the regulatory domain. This preventive external application of high doses of NO˙ by seemingly defenseless wasp eggs represents an evolutionary key innovation that adds a remarkable novel facet to the array of functions of the important biological effector NO˙.

## Introduction

Microbes pose a major threat to the health of all animals and plants. These have responded by evolving a great diversity of defenses including hygienic behaviors [1], antimicrobial chemicals [2-4], complex immune systems [5,6], and defensive symbioses [7,8]. Besides such pathogenic effects, many bacteria and fungi are severe, but often neglected, competitors of animals for nutrients, thus prompting the evolution of mechanisms to preserve food sources [9,10].

Some animals are particularly prone to suffer from microbial attack due to (1) high abundance of potentially harmful microbes in their environment, (2) a microbe-friendly microclimate and/or (3) limited defense mechanisms. The progeny of many insect species develop under warm and humid conditions in the soil, where they are exposed to a high diversity of bacteria and fungi. Moreover, compared to adult insects, immature stages, in particular eggs, have usually reduced abilities to prevent microbial infestation due to, for example, a thin cuticle or an inability to groom [11,12]. The situation is even aggravated when eggs and larvae have to develop on limited amounts of provisions that are susceptible to attack by ubiquitous and fast growing putrefactive bacteria and mold fungi [9,13].

Such hostile conditions prevail in nests of subsocial Hymenoptera like the European beewolf *Philanthus triangulum* (Hymenoptera, Crabronidae). The offspring of these solitary digger wasps develop in subterranean brood cells provisioned by the female wasps with paralyzed honeybee workers (*Apis mellifera*, Apidae, Hymenoptera) [14] (Fig. 1A). The beewolf egg is laid on one of the bees, the larva hatches after three days, feeds on the bees for six to eight days, then spins a cocoon and either emerges the same summer or hibernates. The warm and humid microclimate in the brood cell promotes larval development, but also favors fast growing, highly detrimental fungi [15]. Without any countermeasures the provisions will be completely overgrown by mold fungi within three days (Fig. 1B), and the beewolf larva becomes infested by fungi or starves to death [16,17].

**Figure 1.**
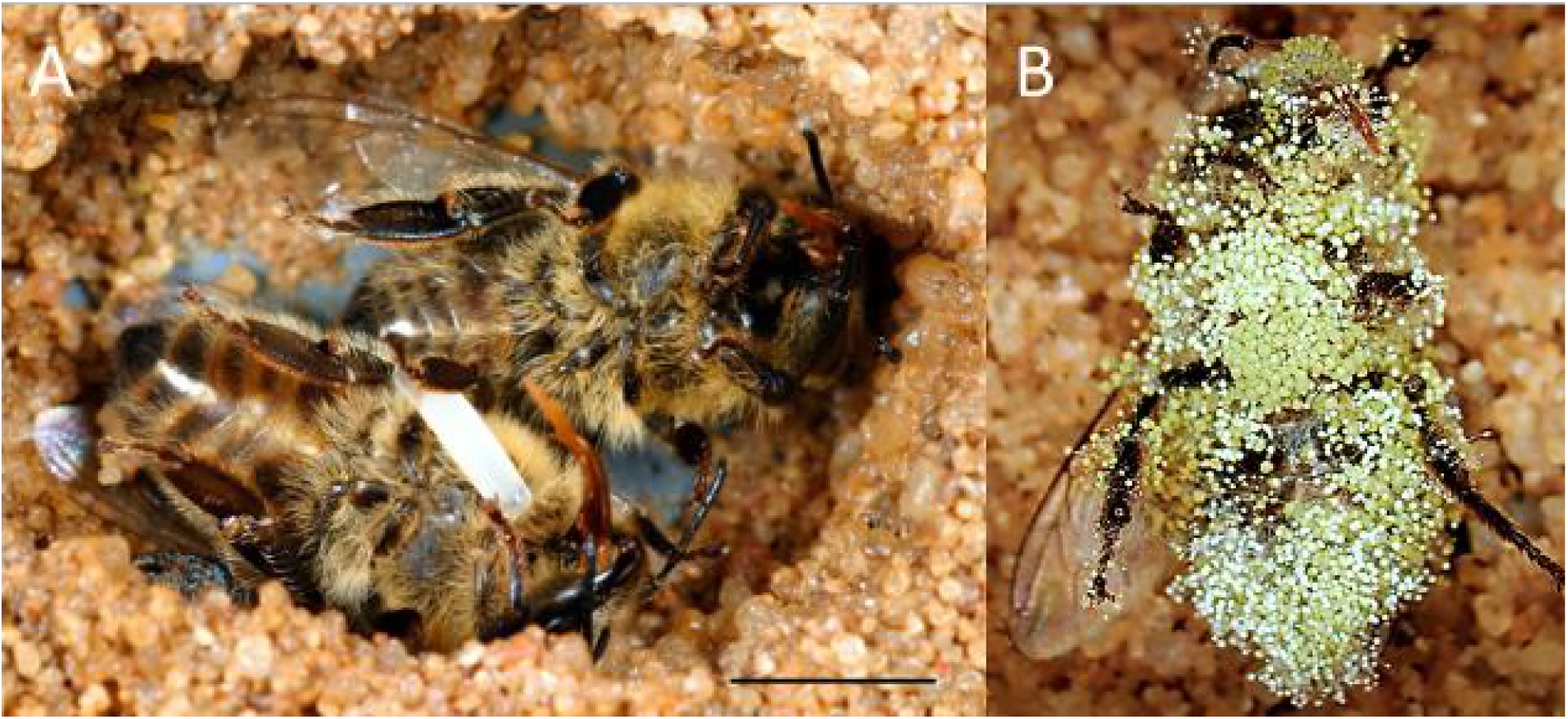
(A) Brood cell of the European beewolf with two bees, one carrying an egg, in an observation cage. (B) Honeybee paralyzed by a beewolf female but immediately removed and kept in an artificial brood cell, heavily overgrown by mold fungi that have already developed conidia. Scale bar = 5mm.

We have previously documented how beewolf females reduce molding of the larval provisions by coating the paralyzed bees with ample amounts of unsaturated hydrocarbons [18]. This embalming changes the physicochemical properties of the preys’ surface causing reduced water condensation on the bees [19]. Due to the deprivation of water, germination and growth of fungi is delayed by two to three days [20]. Despite this considerable effect, when removed from brood cells at least 50% of embalmed bees showed fungus infestation within six days after oviposition [16]. Since in natural brood cells only around 5% of the progeny succumb to mold fungi [16], we searched for an additional antimicrobial defense mechanism.

Here we report on a unique antifungal strategy that is employed by beewolf eggs to defend themselves and their provisions against mold fungi. Employing bioassays we discovered that beewolf eggs emit a strong antifungal agent that we identified as the gaseous radical nitric oxide (NO˙). We characterize the amount and time course of emission and, using histological methods, inhibition assays, and gene expression analysis, we elucidate the biosynthetic pathway of NO˙ in beewolf eggs. To explore the evolutionary background of this remarkable antimicrobial strategy, we sequenced the relevant gene and mRNA. Our findings reveal a novel function of the eminent biological effector NO˙ in providing an extended immune defense to the producer by sanitizing its developmental microenvironment.

## Results

### Emission of an antifungal volatile by beewolf eggs

Thorough examination of beewolf nests in observation cages [21] revealed that within 24 h after oviposition, a conspicuous pungent smell emanated from the eggs. We hypothesized that this smell was due to an antifungal agent. When paralyzed honeybees from completed beewolf brood cells were incubated individually, bees carrying an egg showed significantly delayed fungus growth compared to bees without egg over the period from oviposition to cocoon spinning (Kaplan Meier survival analysis, Breslow test, day 0-11: Chi square = 12, df = 1, p = 0.001; Fig. 2A). This difference was also significant for the period from oviposition to the hatching of the larvae (day 0-3: Chi square = 9.5, df = 1, p = 0.002), suggesting that this effect is not due to possible antifungal mechanisms of the larvae but that it is mediated by the egg. Considering the distinctive odor that emanated from the eggs, we tested whether the antifungal effect is caused by a volatile agent. Two experiments supported this assumption. First, provisioned bees without wasp eggs that were kept in artificial brood cells together with bees carrying an egg (but without physical contact) showed significantly delayed fungal growth compared to control bees that were kept alone (Breslow test, day 0-11: Chi square = 7.6 df = 1, p = 0.006; day 0-3: Chi square = 9.1, df = 1, p = 0.003; Fig. 2B). Second, when one of the most abundant mold species from infested beewolf brood cells, the fast growing *Aspergillus flavus* [15], was exposed to the volatiles presumably emanating from beewolf eggs on nutrient agar for three days, its growth was entirely inhibited, whereas it thrived in controls (binomial test: N = 20, p < 0.001, Fig. 3). In analogous bioassays, five other fungal strains (*A. flavus* strain B, *Mucor circinelloides, Penicillium roqueforti, Candida albicans* and *Trichophyton rubrum*) were similarly inhibited when exposed to volatiles from beewolf eggs (for each strain N = 8, p < 0.01). Notably, when the beewolf larvae were removed from the assays shortly after hatching (three days after oviposition), no fungal growth occurred in the exposed areas during another three days. We conclude that beewolf eggs release a volatile compound with broad spectrum fungicidal properties.

**Figure 2.**
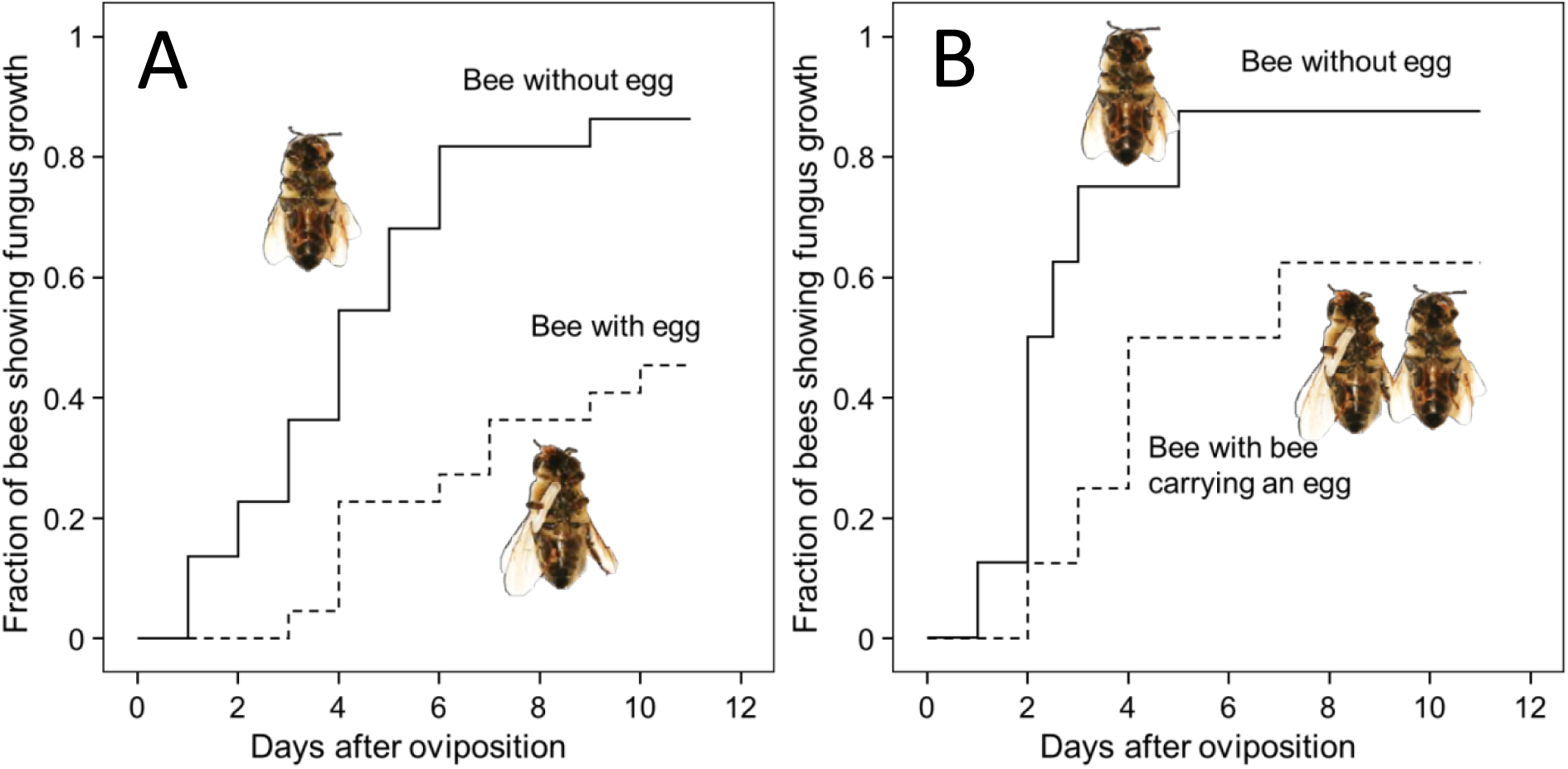
Onset of fungal growth on paralyzed honeybees taken from *Philanthus triangulum* nests and kept in artificial brood cells. The fraction of bees showing first signs of fungal growth is shown as a function of days since oviposition. (A) Honeybees that either carried an egg (dashed line) or not (solid line) (N = 22 each, hazard ratio = 0.43). (B) Honeybees that were either kept alone (solid line) or shared a brood cell with a bee carrying an egg (dashed line) (N = 16 each, hazard ratio = 0.49). Source data file: Fig 2 Source data – effect of egg on fungus growth.xlsx

**Figure 3.**
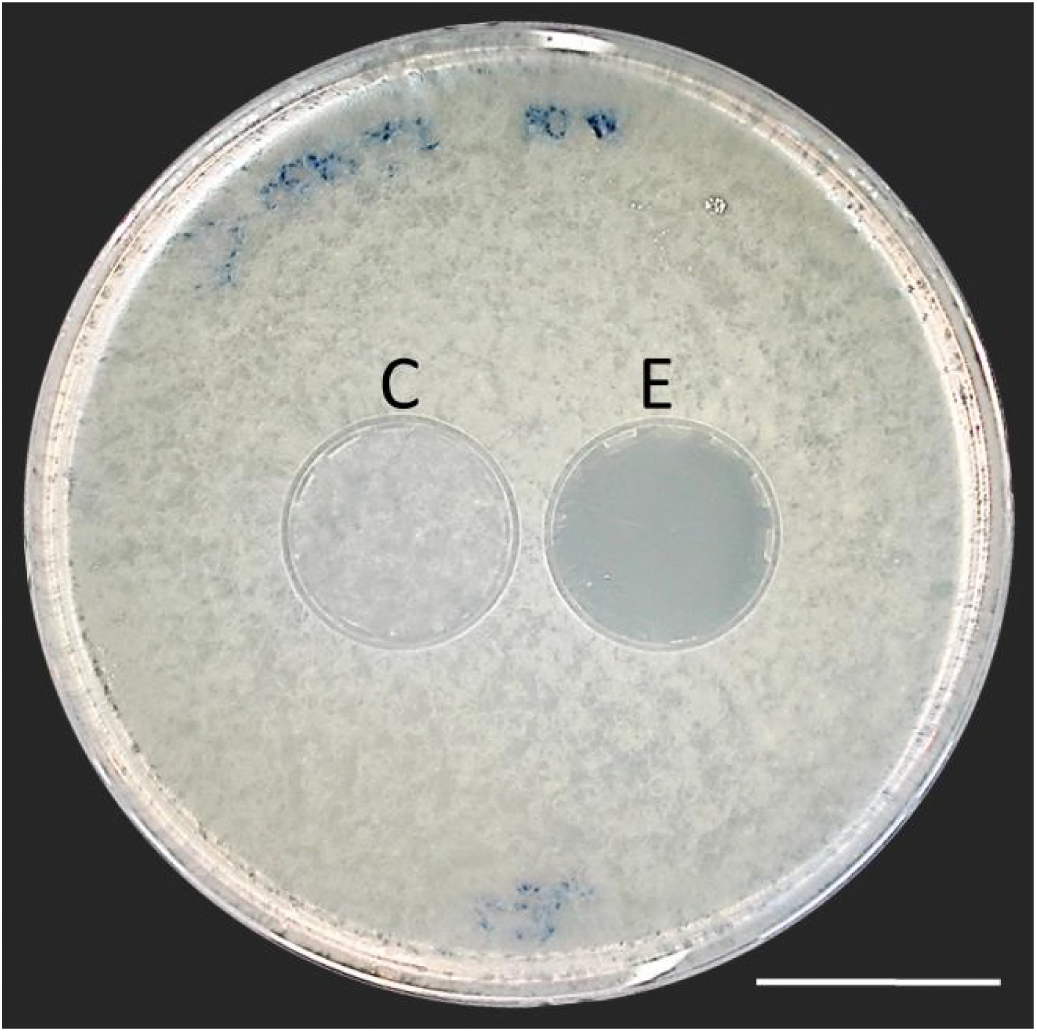
Bioassay demonstrating the inhibitory effect of a beewolf egg against *Aspergillus flavus*. Two areas on the agar were covered by caps of a volume similar to natural beewolf brood cells. One cap, the control (C), was empty, while the experimental cap (E) contained a fresh beewolf egg attached to the ceiling of the cap. The caps were removed and the picture was taken after 24 h of incubation at 25°C. The control area (C) shows dense whitish fungal hyphae similar to the surroundings. However, the area that was exposed to the volatiles from a beewolf egg (E) shows bare agar, indicating that the growth of this aggressive fungus was entirely inhibited. Scale bar = 2.5 cm

### Identification of the antifungal volatile

The odor emanating from the eggs was similar to that of highly reactive oxidants such as chlorine, ozone and nitrogen dioxide [22]. The most likely candidate was the radical nitrogen dioxide (NO_2_˙), because there is a plausible way for it to be generated by wasp eggs: Insect embryos synthesize small amounts of nitric oxide (NO˙) as signaling effectors for developmental processes [23]. If such odorless NO˙ was emitted from the egg, it would spontaneously react with oxygen [24,25] to yield the strong-smelling NO_2_˙. Moreover, belonging to the reactive nitrogen species (RNS), NO˙ and NO_2_˙ show considerable antimycotic activity [26,27] that would explain the observed fungicidal effect of beewolf eggs.

We conducted a series of experiments to determine whether beewolf eggs produce and emit NO˙ and/or NO_2_˙. First, headspace samples of confined beewolf eggs were subjected to the Griess assay, the standard procedure for the specific detection of NO˙ and NO_2_˙ [28]. The emerging red color of the resulting azo dye clearly indicated the presence of NO˙/NO_2_˙. To visualize the emission of NO˙ from beewolf eggs, we sprayed a solution of an NO˙ specific fluorescent probe, Diaminorhodamin-4M AM (DAR4M-AM), onto prey bees carrying freshly laid eggs. The small droplets of the DAR4M-AM solution on the bees showed a clear fluorescence around the egg that increased over several hours (Fig. S1B). No such effect was seen on control bees without eggs (Fig. S1A). Moreover, beewolf eggs injected with the DAR4M-AM solution showed a strong fluorescence that peaked about one day after oviposition (N = 45, Fig. 4A). The same treatment yielded only weak fluorescence in the eggs of two other Hymenoptera (the Emerald cockroach wasp, *Ampulex compressa*, N = 9, and the Red mason bee, *Osmia bicornis*, N = 12; Figs. 4C and D) and in newly hatched beewolf larvae (N = 4, not shown). Autofluorescence of beewolf eggs injected with buffer only (N = 10) was negligible (Fig. 4B). These findings strongly imply that beewolf eggs produce and release NO˙, which spontaneously reacts with oxygen to NO_2_˙ radicals.

**Figure 4.**
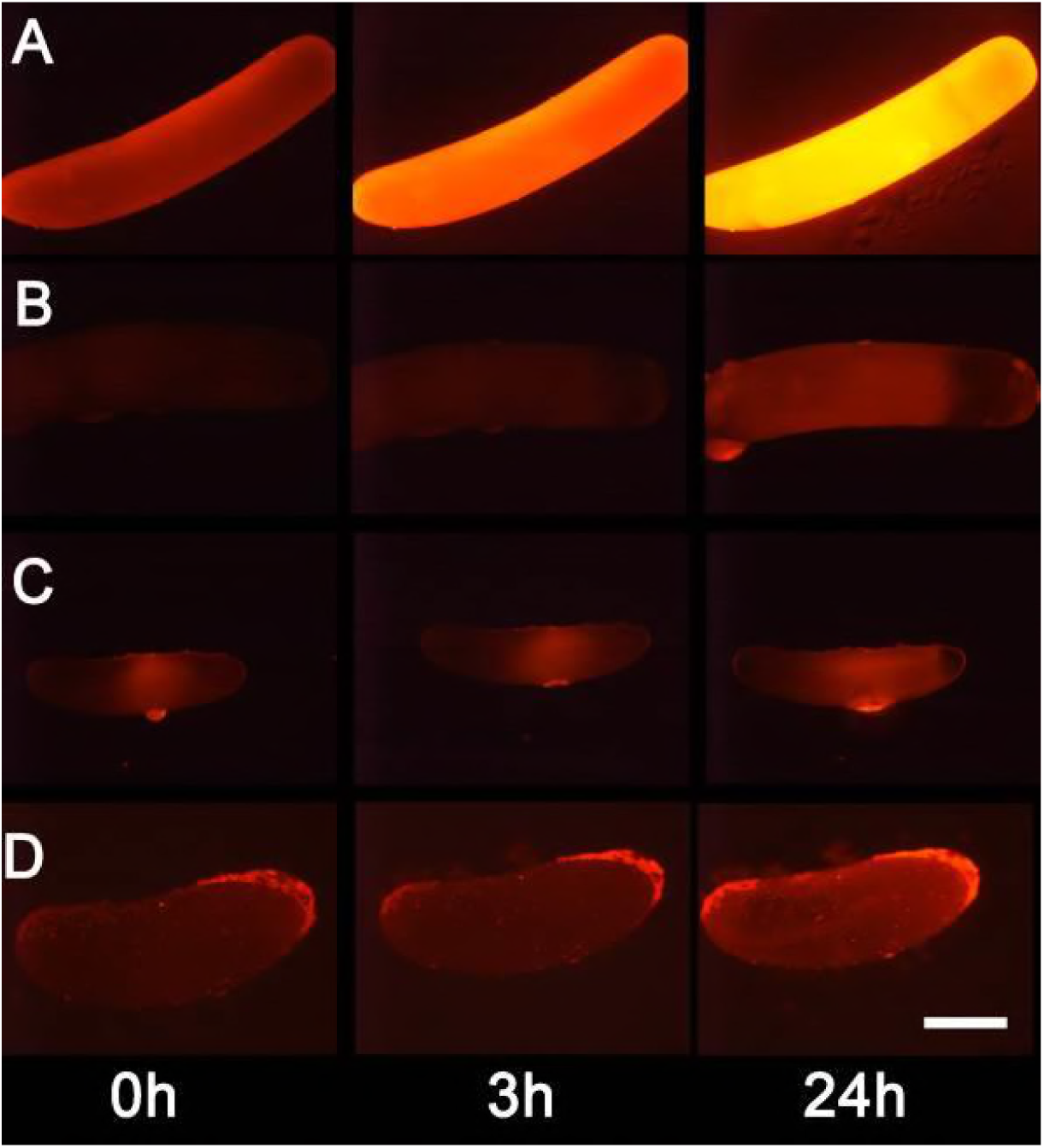
Detection of nitric oxide (NO˙) in beewolf eggs. Newly laid eggs of beewolves, *Philanthus triangulum*, of the cockroach wasp *Ampulex compressa* and of the Red Mason bee, *Osmia bicornis* were injected with the NO˙ sensitive fluorescence probe DAR4M-AM. Control beewolf eggs were injected with phosphate buffer. Images were obtained by fluorescence microscopy 0, 3 and 24 h after injection. Row A: DAR4M-AM injected beewolf egg showing strong increase in fluorescence; B: Buffer-injected control beewolf egg showing the level of autofluorescence; C: DAR4M-AM injected egg of *A. compressa;* D: DAR4M-AM injected egg of *O. bicornis*. Scale bar: 1 mm.

### Amount and time course of NO˙ emission

Using iodometry, we determined that a beewolf egg emits on average 0.25 ± 0.09 µmol NO˙ (N = 233). The rate of NO˙ production was initially very low, but increased to a distinct peak 14-15 h (at 28°C) after oviposition (Fig. 5); around 90 % of NO˙ emission occurred within a two-hour period. Assuming no loss due to reactions or leaking out of the confined space of brood cells (volume 3.2 ± 0.9 cm^3^, N = 250), the nitrogen oxides would accumulate to average concentrations of 1690 ± 680 ppm. The timing of the onset of NO˙ emission was strongly temperature dependent (Fig. S2), with higher temperatures resulting in an earlier NO˙ production (temperature coefficient Q_10_ = 2.74).

**Figure 5.**
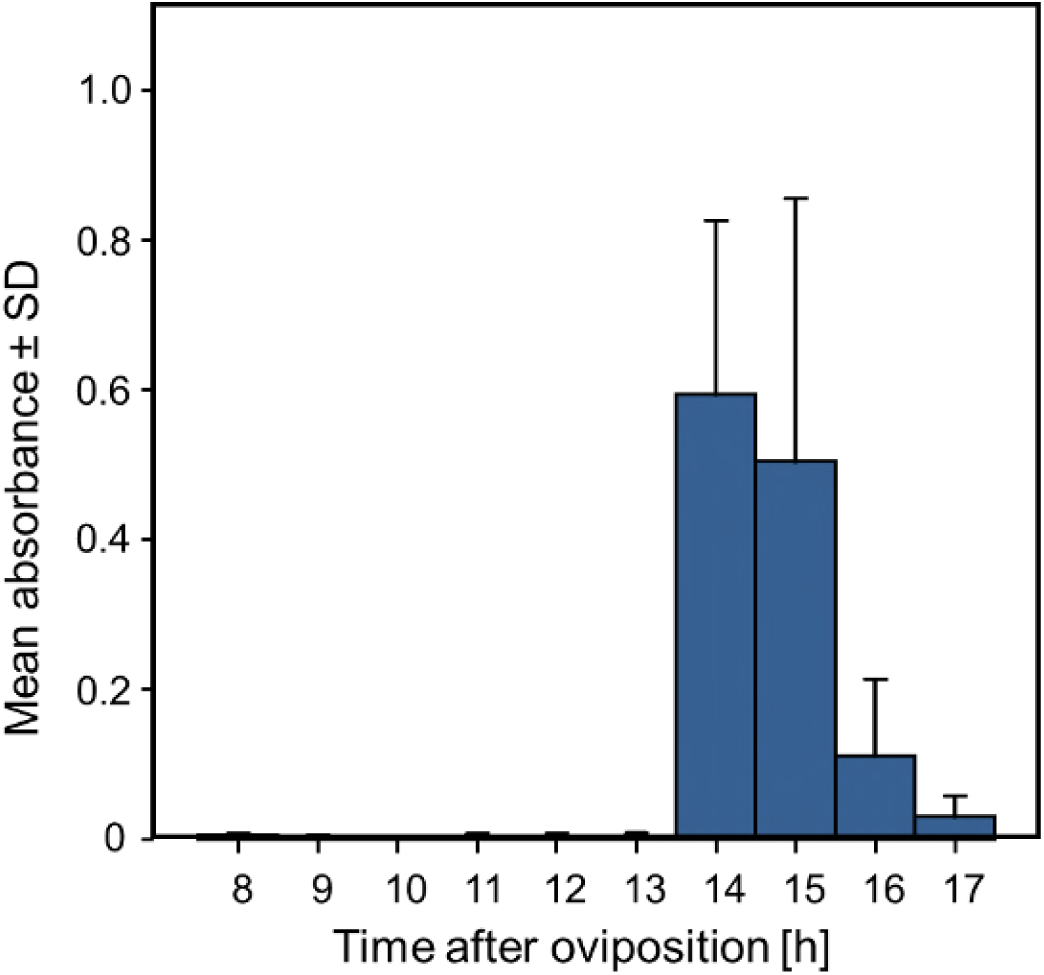
Timing of NO˙ emission from beewolf eggs (N = 4). The photometrically determined absorbance at 590 nm (mean ± SD) is shown as a function of time after oviposition for iodide-starch solutions successively exposed to beewolf eggs for one hour. Source data file: Fig 5 Source data - timing of NO emission.xlsx

**Figure 6.**
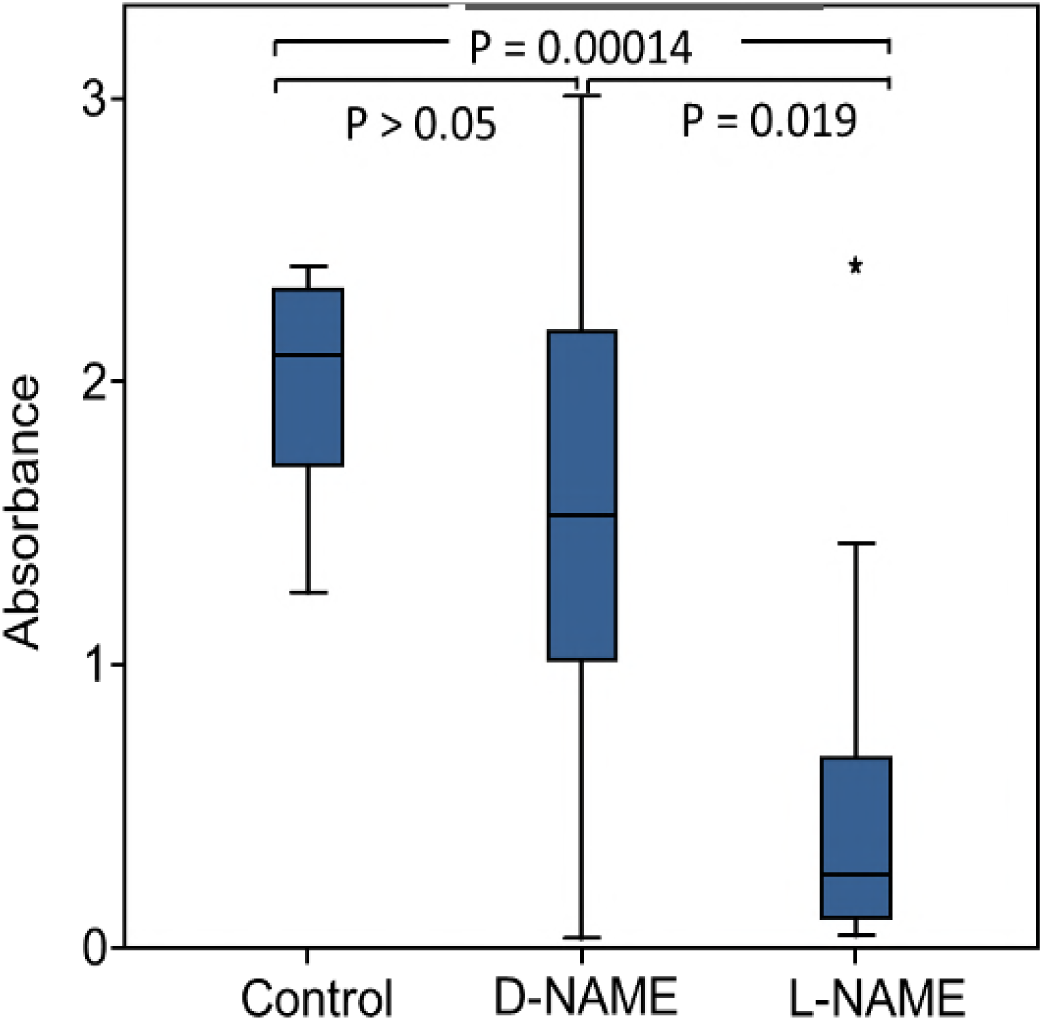
Effect of NOS inhibition on NO˙ production. Amount of NO˙ and/or NO ˙ emanating from non-injected beewolf eggs (control) and those injected with D-NAME (a non-inhibiting enantiomer of L-NAME) or L-NAME (a NOS inhibiting L-arginine analogue). The photometrically determined absorbance at 590 nm is shown for iodide-starch solutions that were exposed for 24 h to the headspace of eggs of the indicated treatment group (shown are median, quartiles and range, * indicates an outlier, included in the analysis). P-values are for Holm-corrected Mann-Whitney U-tests. Source data file: Fig 6 source data – NOS inhibition.xlsx

### Synthesis of NO˙ in beewolf eggs

Eukaryotes synthesize NO˙ from the amino acid L-arginine by the enzyme nitric oxide synthase (NOS) [29] which is highly conserved also in insects [30]. The exceptional level of NO˙ emission of beewolf eggs raised the question of whether they employ the same pathway or have evolved a different mechanism. Using the fixation insensitive nicotinamide-adenine-dinucleotide phosphate (NADPH) -diaphorase assay, we found evidence for NOS activity of embryonic tissue in beewolf eggs (Fig. S3). Moreover, by employing reverse transcription and real time quantitative PCR, we revealed that the temporal expression pattern of NOS-mRNA showed a clear peak around the time of maximum NO˙ emission (Fig. S4). To directly test for the involvement of NOS we injected beewolf eggs with Nω-nitro-L-arginine methylester (L-NAME), a NOS-inhibiting analogue of L-arginine. This treatment caused a significant decrease in NO˙ emission, whereas the non-inhibiting enantiomer D-NAME had no such effect (Fig.6). We therefore conclude that in beewolf eggs, NO˙ is synthesized from L-arginine via NOS.

### The beewolf *NOS*-gene and NOS-mRNA in eggs

In contrast to vertebrates, most invertebrates appear to have only one type of NOS [31,32]. Considering the high level of NO˙ production in beewolf eggs, we hypothesized that beewolves have more than one *NOS* gene or that the NOS responsible for the NO˙ synthesis in beewolf eggs might exhibit considerable changes in enzyme structure compared to the NOS of related species. Sequencing of the *NOS*-gene(s) of *P. triangulum* (*Pt-NOS*) revealed only one *Pt-NOS* copy in the beewolf genome comprising 9.36 kbp with 25 exons (Fig. S5). A phylogenetic analysis of the resulting amino acid sequence revealed a high similarity to the NOS of the closely related bees (Apidae, Fig. S6). However, mRNA sequencing showed that, in contrast to adult beewolves and honeybees, the NOS-mRNA of beewolf eggs (3.72 kbp) lacks exon 14 comprising 144 bp. In the NOS-mRNA of adult beewolves this exon is located between the binding domains for calmodulin and flavin mononucleotide (FMN) (Fig. S5).

## Discussion

Fighting pathogens is of outstanding importance for any organism and has driven the evolution of a great diversity of antimicrobial defenses. Internal immune systems have been extensively documented especially in vertebrates [33-34] but also in insects [35,36], including insect eggs [37]. However, comparatively little is known about external antimicrobial strategies that provide protection for the own body, for the progeny, or for food. There are some reports on the application of antimicrobial secretions on the body surface by adult insects [11,38] or inside a host by larvae of a parasitoid wasp [2]. Carrion beetles preserve the larval food, buried carcasses, by application of antimicrobials [39]. Females of some insect species deposit antimicrobial chemicals [40,41] or antibiotics producing symbiotic bacteria [8, 42] onto their eggs and ant workers can counter microbial infestation of the brood by applying venom [43]. Recently, the employment of volatile antimicrobials by insects as a means of external defense has gathered some interest [4,44.45].

Like other insects that develop in the soil, beewolves are particularly menaced by a diverse and unpredictable range of detrimental microbes. In fact, beewolf progeny and their provisions are under severe threat from fast growing mold fungi [16]. As a consequence, beewolves have evolved at least three very different antimicrobial defenses that provide an effective, coordinated, and enduring protection against a broad spectrum of microbes during the whole development. First, throughout the long period of winter diapause prior to emergence progeny are protected by antibiotics on their cocoons that are produced by symbiotic *Streptomyces* bacteria [7,46,47]. Second, during the early egg and larval stages, molding of the provisions is retarded by an embalming of the honeybees with lipids by the mother wasp [16]. Third, as shown here, the emission of gaseous nitrogen oxide radicals by the beewolf egg might be the most important of the three antimicrobial mechanisms since it takes effect in a very early developmental phase and it results not only in delay of molding but in killing of detrimental fungi in their immediate environment. Thus brood cell fumigation provides beewolf offspring with a decisive initial advantage over the fast growing mold fungi.

The emission of a gaseous agent by beewolf eggs to their confined brood cells is an ideal way to sanitize such intricately structured surfaces as the bodies of honeybees and the rough walls of the brood cell. NO˙ seems to be a most suitable gaseous agent because it can obviously be produced by beewolf eggs in amounts that effectively kill mold fungi in their brood cell. Such volatile sanitation mechanisms that provide a front-line defense against microbes [4] will mostly be inconspicuous and might turn out to be a wider theme in nature.

NO˙ is an ancient biological effector of immense importance in all kinds of organisms ranging from procaryotes to higher plants and animals [29, 48]. Owing to its high diffusibility across biomembranes and specific chemical properties, this gaseous radical plays a crucial role in a multitude of biological processes [29, 48]. In vertebrates, NO˙ is synthesized from L-arginine by three different isoforms of NOS that are encoded by different genes [29, 48]. Low levels of NO (< 1µmol/l) are produced by constitutive NOS (cNOS) isoforms (endothelial eNOS, neuronal nNOS) and have signaling functions, e.g. in neuronal development and in the regulation of vascular tone in vertebrates. Higher NO˙ concentrations (1-10 µmol/l, [49]) are generated by an inducible NOS (iNOS). At such levels NO˙ is highly cytotoxic [49], making it a powerful antimicrobial [26,27], for example in macrophages [29]. However, overproduction of NO˙ due to inflammatory processes [50] or certain diseases (e.g. Alzheimer’s disease, [51]) may cause harmful side-effects [52] and even septic shock [53]. Moreover, NO˙ might affect carcinogenesis and tumor progression in a positive as well as in a negative way [54].

In living tissues, NO˙ is usually removed within seconds by reacting with the heme group of molecules such as oxyhemoglobin [55,56] (very low concentrations may still persist for hours [48]). In air, the autooxidation to NO_2_˙ is comparatively slow so that NO˙ persists (depending on its concentration) for several seconds to minutes [25,56] or even hours [24]. Thus, the NO˙ emitted by beewolf eggs might directly affect fungi, e.g. by damaging DNA [27,57] or by reacting with the heme group of enzymes like cytochrome P450 and cytochrome c oxidase, thus inhibiting these crucial components of the mitochondrial respiratory chain [49,58,59,60]. Yet, most of the antimicrobial activity of NO˙ is attributed to indirect effects via reactive nitrogen species (RNS), in particular nitrogen oxides (NO_2_˙, N_2_O_3_) and peroxynitrite (ONOO^-^, upon reaction with superoxide) [49]. NO_2_˙, has been reported to be severely cytotoxic, e.g. by nitration of tyrosine residues and oxidation of proteins and lipids [26,60].

Even assuming some loss due to reactions or diffusion, the estimated maximum concentration of nitrogen oxides (NO˙ and NO_2_˙) in beewolf brood cells (more than 1500 ppm or 60 µmol/l) vastly exceeds the concentrations in tissues (mostly lower than 0.1 µmol/l [56], 0.85-1.3µmol/l in muscle tissue [61]). The maximum concentration in beewolf brood cells might thus be even higher than what is used in medical applications against multiple drug resistant bacteria (200 ppm NO˙ [62]) or in antifungal treatment of fruit (50-500 ppm NO˙ [63]) and is far beyond permissible exposure limits for humans (e.g. for the USA: 25 ppm for NO˙, 5 ppm for NO_2_˙ [64]).

A beewolf egg of approximately 5 mg emits 0.25 µmol NO˙ within a period of about 2.5 h, or 20.000 µmol/kg*h, a value that is about four orders of magnitude higher than reported baseline levels of NO˙ synthesis in humans (0.15 - ~4.5 µmol/kg*h [65]), rats (0.6-9 µmol/kg*h [66]) and plants (*Arabidopsis thaliana,* 0.36-3 µmol/kg*h [67]), and even considerably higher than in lipopolysaccharide (LPS)-activated macrophages (~800 µmol/kg*h, estimated from [66]). Despite this extremely high rate of NO˙ production in beewolf eggs, the amino acid sequence of the beewolf NOS (*Pt-NOS*) is similar to the closely related bees (Fig. S6), indicating that there was little evolutionary change with regard to the gene itself. Moreover, the structure of the *Pt-NOS* gene is largely homologous to other insects, e.g. *Anopheles stephensi* mosquitoes [68].

However, in contrast to adult beewolves, the NOS-mRNA in beewolf eggs lacks exon 14 (Fig. S5). Such alternative splicing that results in different NOS-mRNAs, including the deletion of exons (but others than in beewolves), has been documented in *A. stephensi* in response to *Plasmodium* infection [69]. Moreover, NOS splice variants may result in organ-specific enzymes in other organisms [29]. Presumably, beewolf eggs produce smaller amounts of another NOS splice variant to support signaling functions in the developing embryo.

In adult beewolves, the exon missing in the NOS-mRNA of eggs is located between the binding domains for calmodulin and FMN. Since calmodulin is believed to be responsible for NOS regulation [70] the deletion of an adjacent part might affect the control of NOS activity in beewolf eggs. Thus, the alternative splicing might enable the production of such large amounts of NO˙. Notably, compared to the cNOS (comprising eNOS, and nNOS) the inducible NOS isoform of vertebrates (iNOS) that generates higher concentrations of NO˙ to combat microbes lacks a section of about 40 amino acids (120 bp) near the FMN domain. Interestingly, this section is thought to be responsible for autoinhibition of the cNOS [71] and its lack enhances NO˙ production by the iNOS. The conspicuous similarity between vertebrate iNOS and the NOS in beewolf eggs with regard to the length of the missing section and its position might suggest a convergent modification to achieve a NOS with high synthetic capacity. Whereas vertebrates have evolved another gene, beewolf eggs appear to accomplish a similar effect by alternative splicing of the mRNA.

The lack of an exon that might circumvent regulation of the NOS and the pattern of *Pt-NOS* expression in the eggs suggest that in beewolf eggs the activity of the enzyme is regulated by gene expression like the NOS in *Plasmodium* infested *A. stephensi* [68] and the iNOS of vertebrates [72,73]. However, in contrast to these cases, in beewolf eggs expression of the Pt-NOS seems not to be induced by immunostimulants but to occur obligatorily at a certain stage in the development of the beewolf embryo.

Even the combined effect of prey embalming and brood cell fumigation does not provide perfect protection as fungus infestation still causes larval mortality in 5% of the brood cells in the field [16]. Some fungal spores might survive under the bees because they were screened against the gas. Another possibility, namely that strains of the ubiquitous mold fungi that are the main causes of molding in beewolf brood cells [15], have evolved resistance against the toxic effects of NO˙/NO_2_˙ seems rather unlikely. There are examples for detoxification of lower concentrations of NO˙ (mainly by scavengers like flavohemoglobins) in different fungi, including species of *Aspergillus* [75,76]. However, the NO˙/NO_2_˙ levels emitted by beewolf eggs are very high and likely affect several very basic biochemical processes, thus making the evolution of an effective resistance unlikely. Moreover, beewolf brood cells are certainly a rare habitat for the ubiquitous mold fungi so that there will be only weak selection for resistance at all.

While brood cell fumigation clearly retards molding of larval provisions, the antimicrobial effect of NO˙ and NO_2_˙ might harm the symbiotic *Streptomyces* bacteria that beewolf females apply to the brood cell prior to egg laying [7,46]. Since the symbiotic bacteria are important for the survival of larvae in the cocoon and are vertically transmitted from beewolf mothers to their daughters [47], a considerable number of symbionts have to survive the brood cell fumigation. Possibly, the symbiotic bacteria are resistant to NO˙/NO_2_˙. However, that the bacteria are applied to the ceiling of the brood cell might be an adaptation that reduces possible negative effects of the nitrogen oxides since these are heavier than air and will accumulate in the lower part of the brood cell. Additionally, the bacteria are embedded in copious amounts of a secretion consisting of mostly unsaturated hydrocarbons [77] that might shield the bacteria from the fumigants. Moreover, antioxidants in the hydrocarbon matrix could detoxify NO˙ and NO_2_˙ and protect the symbiotic *Streptomyces* bacteria.

How could brood cell fumigation with high concentrations of NO˙/NO_2_˙ have evolved? Generally, it has been assumed that the primary purpose of NO˙ was signaling at low concentrations and that the antimicrobial functions of higher concentrations are derived [78]. Assuming a similar scenario for beewolves, small amounts of NO˙ that were originally produced for developmental processes [23] might have accidentally been released into the confines of the subterranean brood cell and slightly affected the germination or growth of fungi by interfering with regulatory processes [29,79]. Given the severe threat posed by microbes, such initial benefits would have caused strong selection for elevated NO˙ emission by the eggs. This would have considerably increased progeny survival and might have allowed ancestral beewolves to nest in an expanded range of habitat types, including nesting sites with high risk of microbial infestation, or to exploit highly susceptible but readily available prey species. Brood cell fumigation with large doses of NO˙ thus represents a key evolutionary innovation. Since NO˙ is used as an antimicrobial in the immune systems of many animals [80], its deployment as an antifungal gas can be viewed as an innate, externalized immune defense of beewolf eggs. Such externalized components of the immune system have recently been recognized as important and possibly widespread antimicrobial measures [38].

The clear benefit of brood cell fumigation, however, is probably accompanied by substantial costs in terms of energy and biochemical resources [31]. NO˙ is synthesized from L-arginine, an amino acid that is an important constituent of many proteins and biochemical pathways [81] and it is an essential amino acid for most insects [82.83] (e.g. phytophagous insects [84], mosquitos [85], aphids [86,87], butterflies [88, 89], true bugs [90], parasitoid wasps [91,92], bees [93,94]). Thus, beewolves have either evolved the capacity to synthesize L-arginine or female beewolves have to provide each egg with sufficient L-arginine for both brood cell fumigation and embryogenesis. Moreover, NO˙ synthesis by NOS requires the cofactors flavin adenine dinucleotide (FAD), FMN, (6R-)5,6,7,8-tetrahydrobiopterin (BH4) and NADPH [95], thus competing with other metabolic pathways in the developing beewolf embryo.

One of the most remarkable aspects of our study is that the embryos inside the egg survive the high concentrations of toxic nitrogen oxides during synthesis and emission as well as after its release to the brood cell. This is all the more surprising since beewolf larvae that were accidentally exposed to the gas emitted by eggs died (Strohm, unpublished observations). The synthesis and emission of such high amounts of NO˙ likely requires a number of concomitant adaptations that protect beewolf embryos against the cytotoxic effects of high concentrations of nitrogen radicals. One possibility is the employment of carrier molecules to transfer NO˙ to the egg shell. In blood sucking hemipterans, for example, nitrophorins carry NO˙ to its release site to dilate blood vessels [96]. The mechanistic basis of NO˙ tolerance of beewolf eggs is of particular interest, since excessive production of NO˙ due to inflammatory processes [97] or certain diseases (e.g. Alzheimer’s disease, [51,52,98,99]) might cause severe pathological complications in humans. Thus, understanding how beewolf eggs avoid the toxic effects of NO˙ might inspire the development of novel medical applications.

Our findings reveal a surprising adaptation in a mass-provisioning digger wasp to cope with the threat of pathogen infestation in the vulnerable egg and larval stages. Sanitizing the brood cell environment by producing high amounts of NO˙ significantly enhances the survival of immatures by reducing fungal growth on their provisions. Given that mass-provisioning and development underground are widespread ecological features among digger wasps and bees and considering the difficulties of detecting volatiles in subterranean nests, such gaseous defenses might be more widespread and as yet underappreciated. In addition to revealing new perspectives on antimicrobial strategies in nature and amplifying the biological significance of NO˙, beewolves offer unique opportunities to elucidate general questions on the evolution and regulation of NOS as well as the production of and resistance to high concentrations of NO˙.

## Methods

### Animals

Beewolf females, *Philanthus triangulum* F. (Apoidea, Crabronidae), were either caught in the field from populations in Franconia (Germany) or were the F1 progeny of such females kept in the laboratory. They were housed in observation cages [21] that provided access to newly completed brood cells. The cages were placed in a room with temperature control (20-22° at night, 25-28°C in the daytime) and were lit for 14 h per day by neon lamps. Honeybees, *Apis mellifera* L. (Apoidea, Apidae), the females’ prey, were caught from hive entrances or from flowers and provided *ad libitum*. Honey was provided *ad libitum* in the flight cage for the nutrition of both honeybees and beewolf females.

To obtain freshly laid eggs, observation cages were checked hourly. Completed brood cells were opened, their length and width was measured using calipers and the egg and/or honeybees were removed and used for the experiments. Brood cell volume was estimated as a prolate spheroid with brood cell length as the major and width as the minor axis. The bees in brood cells had been paralyzed, embalmed with lipids [18], and provisioned by beewolf females. Egg volume was estimated to be 4.1±0.5 mm^3^ (N = 16) by calculating the volume of a cylinder with the respective length and width of an egg (both determined using a stereomicroscope with eyepiece micrometer).

### General experimental procedures

For all experiments beewolf eggs were harvested from brood cells of various females. Eggs were randomly allocated to different treatment groups. Sample sizes refer to independent biological replicates, i.e. each replicate represents a different egg or brood cell – with the exception of quantitative PCR, where several eggs were pooled for one sample (see below). As it is very demanding to obtain beewolf eggs, the availability of eggs of a certain developmental stage was limited. Generally, we used as many eggs as feasible (e.g. for quantitative PCR). For some experiments we decided on a meaningful sample size based on experience from preliminary experiments (e.g. we already knew that inhibition assays with beewolf eggs in Petri dishes were really clear-cut and required only few replicates). Moreover, due to the limited availability of beewolf eggs on a given day, replicates were conducted consecutively over several days.

### Fungus inhibition assays

To test whether the time course of fungus growth on bees differed between those carrying an egg and those without egg, we used brood cells (N = 22) that had been provisioned with two bees. We placed each bee individually into an artificial brood cell of natural shape and volume in sand-filled Petri dishes (diameter 10 cm) and with moisture levels similar to natural conditions. Petri dishes were placed in a climate chamber at 25°C in the dark. Bees were carefully checked visually every 24 h for fungus growth without opening the Petri dishes. First signs of fungus infestation (hyphae) were recorded. The experiment was terminated after eleven days since all larvae had finished feeding and spun a cocoon by then. Since these are time event data, we used survival analysis (Kaplan Meier, Breslowe test, SPSS Statistics 21) to compare the timing of fungus infestation of the bees with and without an egg. Larvae hatched on the third day after oviposition and started to feed on the bee. There was no evidence that hatched larvae were able to prevent fungus growth on the bee they occupied or others in the brood cell. However, to take a possible effect of the larva on the experimental bee into account, we carried out the analysis not only over the whole period from oviposition until the larvae spun into a cocoon (11 days) but also for the period from oviposition to the hatching of larvae (3 days). A significant difference already until the third day indicates that this effect was associated with the egg.

We examined whether beewolf eggs emit a volatile antimicrobial by conducting two experiments. For the first test, we used brood cells (N = 16) that contained three bees. The bees were transferred to artificial brood cells in sand-filled Petri dishes as described above. The bee with the egg and one of the bees without egg (the experimental bee) were placed together in the same artificial brood cell but without physical contact. The other bee without egg (the control) was kept alone in another artificial brood cell (in another Petri dish). We monitored the timing of fungus infestation, as described above. We used survival analysis as described above. Again, to take an (unlikely) effect of the larva into account, we also carried out the analysis for the period from oviposition until the larvae hatched (day 3 after oviposition). A significant difference already until day three could only be caused by volatiles emanating from the egg.

For the second assay, we exposed conidiospores of a diverse spectrum of fungi to the volatiles emanating from beewolf eggs. Petri dishes (10 cm) containing culture medium (malt extract agar or Sabouraud-agar [100]) were inoculated with conidia from different fungal strains (*Aspergillus flavus* strain A, Trichocomaceae, that was isolated from infested beewolf brood cells, [15], N = 20; *A. flavus* strain B, *Mucor circinelloides,* Mucoraceae; *Penicillium roquefortii,* Trichocomaceae; *Candida albicans,* Saccharomycetaceae; *Trichophyton rubrum*, Arthrodermataceae; N = 8 for all the latter strains and species; these were kindly provided by the Department of Hygiene and Microbiology of the Würzburg University Hospital). Conidiospores were harvested by sampling mature fungus colonies that were reared from stock cultures. A suspension of the conidia in sterile water was evenly distributed on the Petri dishes to obtain uniform growth of fungi. To recreate the concentrations of potential antibiotic volatiles in the brood cell, we used small plastic caps (3 ml, about the size of a brood cell) to confine test areas on the agar. Freshly laid eggs were placed singly on the bottom of a cap where they readily attached due to their natural stickiness. Each cap was then placed on a freshly inoculated Petri dish so that the agar under the cap was not in contact with the egg but was exposed to volatiles that emanated from the egg. An empty cap was placed on the same Petri dish as a control. The Petri dishes were incubated in a dark climate chamber at 25°C. Fungus growth under the experimental and control caps was recorded after 24, 48 and 72 h. After 72 h the caps with the hatched larvae were removed, and fungal growth was further recorded after another 24, 48 and 72 h. Since there was either no fungal growth or substantial growth (Fig. 3), the experimental and control areas were compared using binomial tests (software PAST [101]). The qualitative results for all other observation times were identical to those after 24 h.

### Identification of the antimicrobial volatile

We hypothesized that nitric oxide (NO˙) and its main reaction product with oxygen, nitrogen dioxide (NO_2_˙), were the most likely compounds emanating from beewolf eggs. The standard test for the detection of NO˙ and NO_2_˙ employs the Griess reaction. We used a solution of sulfanilic acid and N-(1-naphthyl)-ethylenediamine (Spectroquant Nitrite Test, Merck, Germany, according to the manufacturer’s instructions). The Griess reagent specifically reacts with the nitrite anion (NO_2_^-^) to form a distinctive red azo dye [102]. NO˙ reacts with water to form nitrous acid (HNO_2_) and can thus be directly verified by the Griess reaction. NO_2_˙, however, disproportionates in water into nitrous acid and nitric acid (HNO_3_) and the latter must be reduced to nitrous acid to react with the Griess reagent. Freshly laid beewolf eggs (collected within 2 h after oviposition, N = 11) were placed in the lid of a 1.5 ml reaction tube where they readily attached due to their natural stickiness. Tubes without eggs (N = 11) were used as controls. Then 1 mL of the Griess test solution was added to the tube. For another sample (N = 15) the nitrate, which might be present in the solution, was reduced to nitrite by placing a glass fiber filter disc with small amounts of zinc powder [103] on the surface of the solution. The same setting without an egg was used as control (N = 15). The tubes were incubated at 25°C for 24 h, and the occurrence of the red coloration was examined both visually and with a photometer (at 520 nm, Nanophotometer, Implen, Germany). The samples with and without nitrate reduction showed the same results.

NO˙ can also be detected by specific fluorescent probes. In particular, diaminorhodamin-4M AM (DAR4M-AM), a cell permeable, photostable fluorescent dye, has a high sensitivity and specificity for NO˙ [104]). A DAR4M-AM (Alexis Biochemicals, USA) solution was prepared according to the supplier’s instructions (10µmol/l in 0.1mol/l phosphate buffer, pH 7.4). To verify and to visualize the emission of NO˙ from the egg, paralyzed honeybees either with freshly laid eggs (N = 8) or controls without eggs (N = 8) were sprayed with the DAR4M-AM solution using a nebulizer (the egg itself was screened from droplets during spraying) and kept in the dark (at 25°C in artificial brood cells as described above). After 20 h, the bees were examined under a fluorescence microscope (Axiophot II, Zeiss, Germany, filter set 43: excitation 520-570 nm, emission 535-675 nm) and digital photos were taken (Nikon DS-2 Mv, Nikon Japan) at constant exposure times, to allow comparison of fluorescence intensity. Due to the size of the bees, several pictures had to be taken in the X,Y plane as well as along the Z axis. Pictures along the z-axis were stacked using the software Combine-ZP (www.hadleyweb.pwp.blueyonder.co.uk). Then these stacks were stitched using Photoshop Elements 5 (PSE5, Adobe Systems Inc. USA). Since small peripheral background parts within the frame of the stacked and stitched picture were “empty” these parts were filled with other background parts by using the clone stamp tool. Images were corrected for contrast and sharpness using PSE5 with identical settings for experimental and control specimens.

DAR4M-AM can also be used to detect NO˙ in tissues. Aliquots of 0.1-0.5 µl of the DAR4M-AM solution (see above) were injected into beewolf eggs (within 1 h after oviposition, N = 64, in N = 45 eggs the embryo survived and developed) with a custom made microinjector equipped with glass capillaries (Eppendorf Femtotips II, Eppendorf, Germany) under microscopic control. The eggs were kept in dark chambers (as above), and fluorescence was observed directly after injection and 1, 3, 5, 24, 48 and 72 h later. Control eggs injected with buffer only (N = 10) were monitored in the same way to assess autofluorescence. For comparison, eggs of two other Hymenoptera (*Osmia bicornis*, Apoidea, Megachilidae, N = 12, and *Ampulex compressa*, Apoidea, Ampulicidae, N = 9; eggs from both species were obtained from our own laboratory populations) as well as freshly hatched beewolf larvae (N = 4) were injected with the DAR4M-AM solution and monitored in the same way. Fluorescence was examined under a fluorescence microscope and documented with a digital camera as described above. Contrast and sharpness of the images were optimized using Photoshop Elements 5 (Adobe, USA) with identical settings for all specimens.

### Quantification, time course and temperature dependence of NO˙ production

Iodometry provides a simple but sensitive, reliable and precise method to quantify strong oxidants. To assess the amount of emitted nitrogen oxides, we placed freshly laid eggs (N = 233) individually into the lid of 1.5 ml reaction tubes where they readily attached due to their natural stickiness. Then 1 ml of a potassium iodide-starch solution (containing 1% KI and 1% soluble starch in distilled water) was added, the reaction tube was closed and kept for 24 h at 28°C in a dark climate chamber. Oxidation of iodide results in iodine that forms a blue complex with starch [103]. The degree of coloration was quantified by measuring the absorbance at 590 nm in a spectrophotometer (Uvikon 860, Kontron, Germany). To assess the absolute amount of the oxidant, the solutions were subsequently calibrated by titration with a reference solution of sodium thiosulfate (concentration: 0.001 M; Merck, Germany) until the blue color of the iodine-starch complex disappeared.

To establish the time course of gas production, individual beewolf eggs (N = 4) were transferred within 1 h after oviposition into the lid of reaction tubes and kept in a dark climate chamber at 28°C. Every hour, the cap with the egg was transferred to another reaction tube with fresh iodide-starch solution. Immediately after removal of the egg from a reaction tube, absorbance of the solution was measured at 590 nm as described above.

To investigate the temperature dependence of gas production, tubes with a newly laid egg and iodide-starch solution (as described above, N = 33 in total) were placed in a rack (with white background) inside a climate chamber and incubated at seven different constant temperatures (20, 22.5, 24, 25.5, 27, 28.5 and 30°C). The time course of coloration of the iodide-starch solution was recorded using a digital camera (Canon EOS 20D, Canon, Japan) programmed to take pictures at 30 min intervals. The onset of gas production could be easily determined since the color of the solution turned from clear to dark blue from one picture to the next, i.e. within a 30 min interval. A quadratic regression curve was fitted to the data (SPSS Statistics 21) and the Q_10_ value for the temperature dependence was estimated.

### Detection of NOS activity in egg tissue

To detect NOS activity in the egg tissue, we used fixation-insensitive NADPH diaphorase staining with nitroblue tetrazolium [105,106]. Eggs were fixed in PBS containing 4% paraformaldehyde for 2 h at 4°C, followed by cryoprotection in PBS with 12% sucrose for 20 h. The tissue was soaked in Tissue Tec (Sakura Finetek, Netherlands) for 30 min, frozen, and 10 µm sections were cut on a cryostat microtome (CM3000, Leica, Germany). The sections were incubated for 60 min at 30°C with 50 mmol/l Tris-HCI, pH 7.8, 0.1% Triton X-100, and 0.2 mmol/l nitroblue tetrazolium chloride in the presence or absence (each N = 5) of 0.2 mmol/l β-NADPH to demonstrate fixation-insensitive NADPH diaphorase activity. The sections were dehydrated, mounted with Depex (Serva, Germany) and observed under a compound microscope (Zeiss Axiophot II). Photos were taken with a digital camera (Nikon DS-2 Mv). To cover the whole egg, two pictures had to be stitched (Photoshop Elements 5, Adobe USA) and contrast and sharpness were optimized.

### Phenology of NOS gene expression

If NOS is responsible for NO˙ production in beewolf eggs, the time pattern of *NOS* gene expression should largely resemble the time course of NO˙ production by showing a pronounced peak several hours after egg laying (the timing of the peak depending on temperature). We used reverse transcription and real time quantitative PCR to quantify the NOS mRNA in beewolf eggs at different times after oviposition. Since the amount of mRNA that could be obtained from single eggs was insufficient to get reproducible results, we pooled 10 eggs in one trial and 24 eggs in a second trial for each of four different time intervals after oviposition (4-5, 9-10, 14-15 and 19-20 h after oviposition, all kept at 25°C) as well as the respective number of freshly hatched larvae. The eggs and larvae were removed from the brood cells at the specified times, shock frozen with liquid nitrogen and stored at −80°C. The RNA of each sample was extracted using the peqGOLD total RNA Kit (Peqlab, Germany) according to the supplier’s instructions and eluted with 20 µL RNase free water. An aliquot of 3 µL of the RNA was digested with DNaseI (Fermentas, Lithuania) and transcribed into cDNA with BioScript (Bioline, Germany) using an Oligo-dT primer (Fermentas, Lithuania) in a final volume of 20 µL. As a reference for basic levels of gene expression during the experimental period, mRNA of the housekeeping gene β-actin was quantified and the ratio of NOS/β-actin mRNA was calculated for each sample.

For quantitative PCR, we established new primers for both the NOS and β-actin genes of *P. triangulum* (based on the complete *NOS* sequences, see below) (NOS_qPCR_F1 & R4; Actin_qPCR_F1 & R1, Table S1). All primers were intron-overlapping to avoid the measurement of contaminating genomic DNA. The NOS and actin primers amplified fragments of 312bp and 321bp, respectively. The qPCRs were performed on an Eppendorf Realplex cycler (Eppendorf, Germany) in a final volume of 25 µL, containing 1 µL of template cDNA (1 µL of the 20 µL RT reaction mix), 2.5 µL of each primer (10 pmol/l) and 12.5 µL of SYBR Green Mix (SensiMixPlus SYBR Mit, Quantace, UK). Standard curves were established by using 10^-9^ – 10^-3^ ng of PCR products as template. A NanoDrop TM1000 spectrophotometer (Peqlab, Germany) was used to measure DNA concentrations of the templates for the standard curves. PCR conditions were as follows: 95°C for 5 min, followed by 50 cycles of 56°C (β-actin) or 65°C (NOS) for 60 s, 72°C for 60 s and 95°C for 60s. Then a melting curve analysis was performed by increasing the temperature from 60°C to 95°C within 20 min. Based on the standard curves, the amount of NOS and β-actin template and their ratio was calculated.

### NOS inhibition assay

To verify the role of NOS in NO˙ production by beewolf eggs, we used an inhibition assay [107]. Since L-arginine is the substrate for NO˙ production by NOS, we injected either an inhibiting L-arginine analogue or, for controls, a non-inhibiting enantiomer into freshly laid beewolf eggs. Chemicals were dissolved in 0.1 mol/l phosphate buffer pH 7.4. Using a microinjector (see above) eggs were injected with about 0.2 µl of 1.5 mol/l solutions of (1) the competitive inhibitor Nω-nitro-L-arginine methylester (L-NAME, Sigma-Aldrich, USA) (experimental group, N = 14), or (2) the non-inhibiting Nω-nitro-D-arginine methylester (D-NAME, Sigma-Aldrich, USA) (control group 1, N = 9) or (3) not injected at all (N = 14, control group 2). Each egg of the three groups was placed individually in the lid of a reaction tube with an iodide-starch solution as described above and incubated for 24 h at 28°C. Then NO˙ production was assessed by measuring absorbance of the solution with a photometer (Implen Nanophotometer) at 590 nm. Statistical comparison of the groups was conducted using Mann-Whitney U-tests with correction after Holm [108] (SPSS Statistics 18).

### Sequencing of the *P. triangulum* nitric oxide synthase gene (*Pt-NOS*) and mRNA

DNA was extracted from female beewolf heads with the Epicentre MasterPure Complete DNA and RNA Purification kit (Epicentre, USA) according to the manufacturer’s guidelines for tissue extraction. Eggs for RNA extraction were kept at a temperature of 27.5°C (range 26-29°C), collected 14-15 h after oviposition, immediately frozen in liquid nitrogen and stored at −70°C until RNA extraction. Twenty eggs were pooled for extraction and homogenized by repeatedly pipetting in lysis buffer of the PeqGOLD Total RNA kit (Peqlab, Germany). Samples were processed according to the kit manual and frozen at −70°C. For the full transcriptome sequencing (to obtain the 5’ terminal region) RNA was extracted from the antennae of eight frozen female beewolves according to manufacturer’s protocol 1 of the innuPrep RNA Mini Kit (Analytik Jena, Germany).

Most of the beewolf *NOS* gene was amplified and sequenced by primer walking. Sequencing reactions were performed by a commercial service (Seqlab, Germany). Four degenerate primers (NOS860fwd2, NOS1571rev1, NOS_seq_F1_deg, and NOS_seq_R1_deg) were designed (Table S1) based on published *NOS* sequences of *Drosophila melanogaster* (U25117.1), *Apis mellifera* (AB204558.1), *Anopheles stephensi* (AH007775.1), *Rhodnius prolixus* (U59389.1), *Manduca sexta* (AF062749.1) and *Nasonia vitripennis* (NM_001168232.1). First, the central region (~700bp, between NOS860fwd2 & NOS1571rev1) was amplified and sequenced. Based on this sequence, we designed a pair of *P. triangulum* specific primers (NOS_qPCR_F2 and NOS_qPCR_R2, Table S1). Using one specific central and one degenerate terminal primer (NOS_seq_F1_deg and NOS_seq_R1_deg, Table S1), respectively, fragments of 4-5 kb were amplified and sequenced by primer walking, which yielded the central 9.5 kb of the *NOS* gene.

Fragments larger than 2 kb were amplified with the PeqGOLD Mid-Range PCR System on a thermocycler (TGradient, Biometra, Germany). Reaction volumes of 12.5 µL contained 1 µL DNA template, 50 mmol/l Tris-HCl (pH 9.1), 14 mmol/l (NH_4_)_2_SO_4_, 1.75 mmol/l MgCl_2_, 350 mmol/l of each dNTP, 400 mmol/l of each primer and 0.5 U ‘MidRange PCR’ enzyme mix. An initial 3 min melting step at 94°C was followed by 35 cycles of 0.5 min at 94°C, 0.5 min at 58°C and 3 min+20 sec per cycle at 68°C and a final extension time of 20 min at 68°C.

Fragments up to 2 kb were amplified using the PeqGOLD Taq. Reaction volumes of 12.5 µL contained 1 µL of DNA template, 50 mmol/l Tris-HCl pH 9.1, 14 mmol/l (NH_4_)_2_SO_4_, 3 mmol/l MgCl_2_, 240 µmol of each dNTP, 800 nmol/l of each primer and 0.5 U Taq. An initial 3 min melting step at 95°C was followed by 35 cycles of 1 min at 95°C, 1 min at 60°C and 2 min at 72°C and a final extension time of 3 min at 72°C.

The 3’ terminus was sequenced following the 3’ RACE protocol [109]. Briefly, cDNA was generated by reverse transcription with a poly-T primer. Before reverse transcription, co-extracted DNA was digested using DNaseI (New England Biolabs, UK). The DNA digestion mix contained 1 mmol/l Tris-HCl, 0.25 mmol/l MgCl_2_ and 1 mmol/l CaCl_2_ and 0.4 U DNaseI. DNA was digested for 10 min at 37°C, followed by DNase inactivation for 10 min at 75°C. The final reverse transcription mix contained 25 mmol/l KCL, 10 mmol/l Tris-HCl, 0,6 mmol/l MgCl_2_, 2 mmol/l DTT, 4 µmol poly-T or gene specific primer, 0.5 mmol/l of each dNTP and 200 U of BioSkript Moloney Murine Leukaemia Virus reverse transcriptase (Bioline, Germany). The entire digestion mixture was incubated with the primer for 5 min at 70°C to enable primer annealing, then cooled on ice. Reverse transcription was carried out for 1 h at 42°C and the enzyme was subsequently inactivated for 10 min at 70°C. The cDNA including the 3’ terminal region was amplified with the specific primer NOS_seq_3-F3 and a ‘poly-T adapter primer’, i.e. a polyT primer to which a specific adapter sequence was added [109] (Table S1). Subsequently, a nested PCR was performed using a second specific primer (NOS_seq_3-F6) and a primer that contained only the specific adapter sequence of the ‘polyT adapter primer’ to increase PCR specificity (Table S1, same PCR conditions as above).

The 5’ terminal region of 200 bp was obtained from a full transcriptome sequencing approach of female antennae, which covered the full-length NOS mRNA sequence. RNA sequencing was performed by a commercial service provider (Fasteris, Switzerland), using the HiSeq TM2000 Sequencing System (Illumina, USA) with 100 bp single reads, on 5 μg total RNA isolated from female *P. triangulum* antennae. CLC Genomics Workbench was used for sequence assembly of the resulting 75 million reads. Reads were quality-trimmed with standard settings and subsequently assembled using the following CLC parameters: nucleotide mismatch cost = 2; insertion cost = 2; deletion cost = 2; length fraction = 0.3; similarity = 0.9. Conflicts among the individual bases were resolved by voting for the base with highest frequency. Contigs shorter than 250 bp were discarded.

To sequence the entire NOS transcript from eggs, cDNA was generated by reverse transcription with a poly-T primer and additionally a specific, central NOS_RT_R1 primer, followed by PCR amplification using various primer combinations to cover the whole transcript sequence (Table S1). Additionally, the sequence of the 5’ terminal region was confirmed by RT-PCR of mRNA from *P. triangulum* eggs, using primers NOS_seq_5-F6 and NOS_seq_5-R3 (Table S1) and subsequent sequencing.

Even though we used a large number of primers to cover the gene, we did not find sections with signals for two different bases at the same site. Thus we infer that there is only one *NOS* gene in the *P. triangulum* genome, as in most invertebrates [110]. In addition, the transcriptome dataset did not reveal any other transcript that was annotated as nitric oxide synthase.

The GenBank accession numbers for the *P. triangulum NOS* (*Pt-NOS*) gene sequence is: KJ425525, for the NOS mRNA of *P. triangulum* eggs: KJ425526, and for the NOS mRNA in *P. triangulum* female antennae: KJ425527.

### Phylogenetic analysis of *NOS* gene sequences

NOS coding sequences of 23 insect species from five orders were acquired from the NCBI database. Along with the *P. triangulum NOS* sequence, these were translated and aligned using Geneious (Version 6.0.5, created by Biomatters, Geneious, New Zealand). The highly variable 5’ end was trimmed. An approximately-maximum-likelihood tree was created with FastTree [111, 112]. Local support values were estimated with the Shimodaira-Hasegawa test based on 1,000 samples without re-optimizing the branch lengths for the resampled alignments [111]. Bayesian estimates were made with the program MrBayes 3.1.2 [113-115). The MCMC analysis was conducted under a mixed amino acid rate model (prset aamodelpr= mixed). After 1,000,000 generations, with trees sampled every 1000 generations, the standard deviation of split frequencies was consistently lower than 0.01. We discarded the first 100 of the sampled trees (10% burn-in) and computed a 50% majority rule consensus tree with posterior probability values for every node. The trees estimated by both methods were nearly identical, so they were combined into a single figure.

## Acknowledgements

We are grateful to Wilhelm Boland for advice, to Nathalie Moske for technical assistance, to Stephan Schneuwly for providing the microinjection device and, in particular, to Jon Seger and Jeremy Field for valuable comments on earlier drafts of the manuscript.

## Author Contributions

Conceptualization ES

Methodology ES TE MK GH JR

Formal Analysis ES TE

Investigation ES TE GH JR MK

Resources ES JR

Writing – Original Draft Preparation ES TE

Writing – Review & Editing ES TE GH MK JR

Visualization ES TE

Supervision ES

Funding Acquisition ES

## Supporting information

### SI Figures

**Figure S1.**
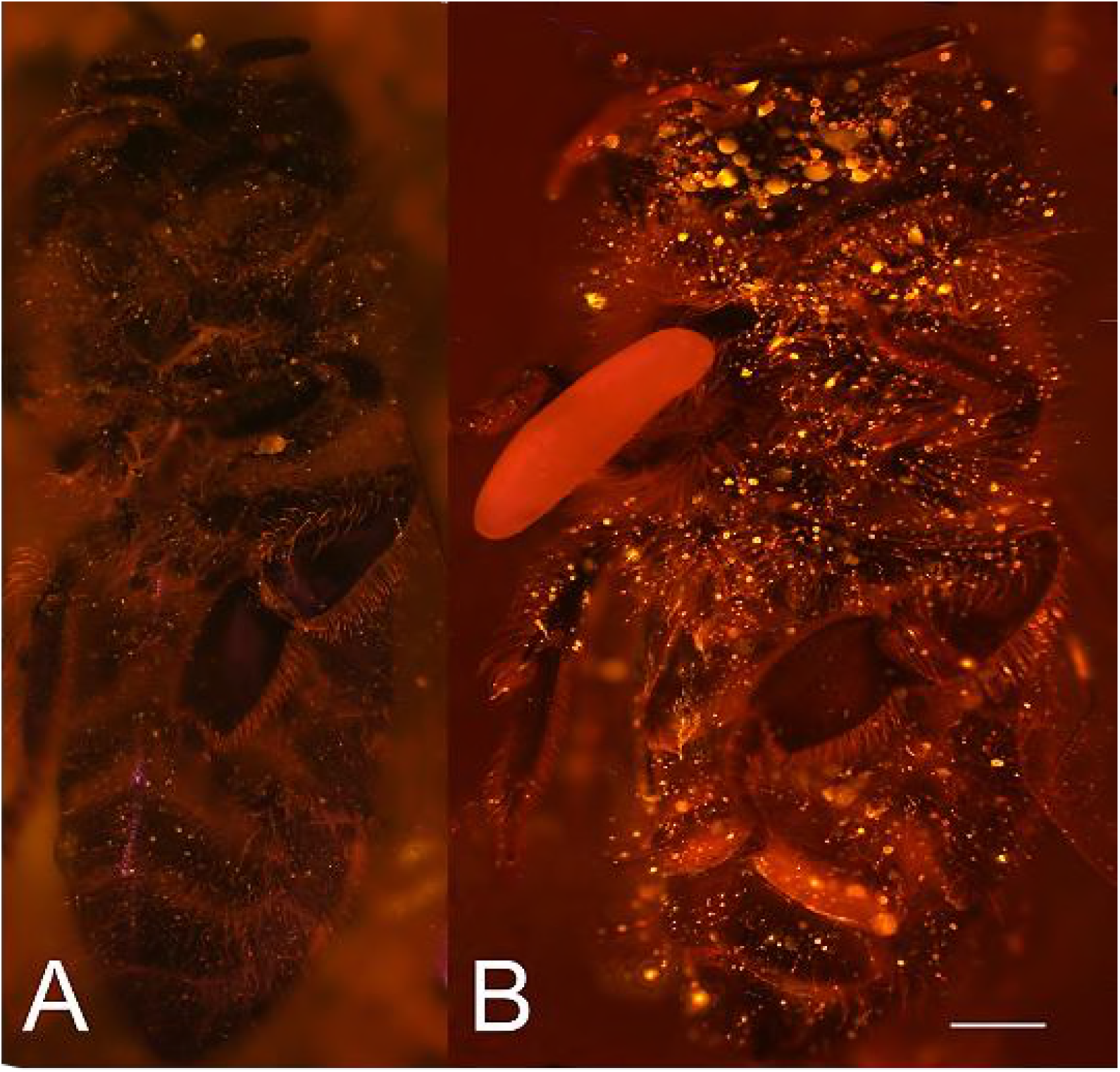
Visualization of NO˙ emission by beewolf eggs using fluorescence imaging. (A) Honeybee from a brood cell without an egg and (B) honeybee with egg. Both bees were sprayed with a solution of the NO˙ specific fluorescence probe DAR4M-AM. Only the droplets on the bee with the egg (B) show a bright yellow and orange fluorescence indicating the presence of NO˙. Image is a composite of multiple pictures of the x/y plane and z-axis. Scale bar = 1 mm.

**Figure S2.**
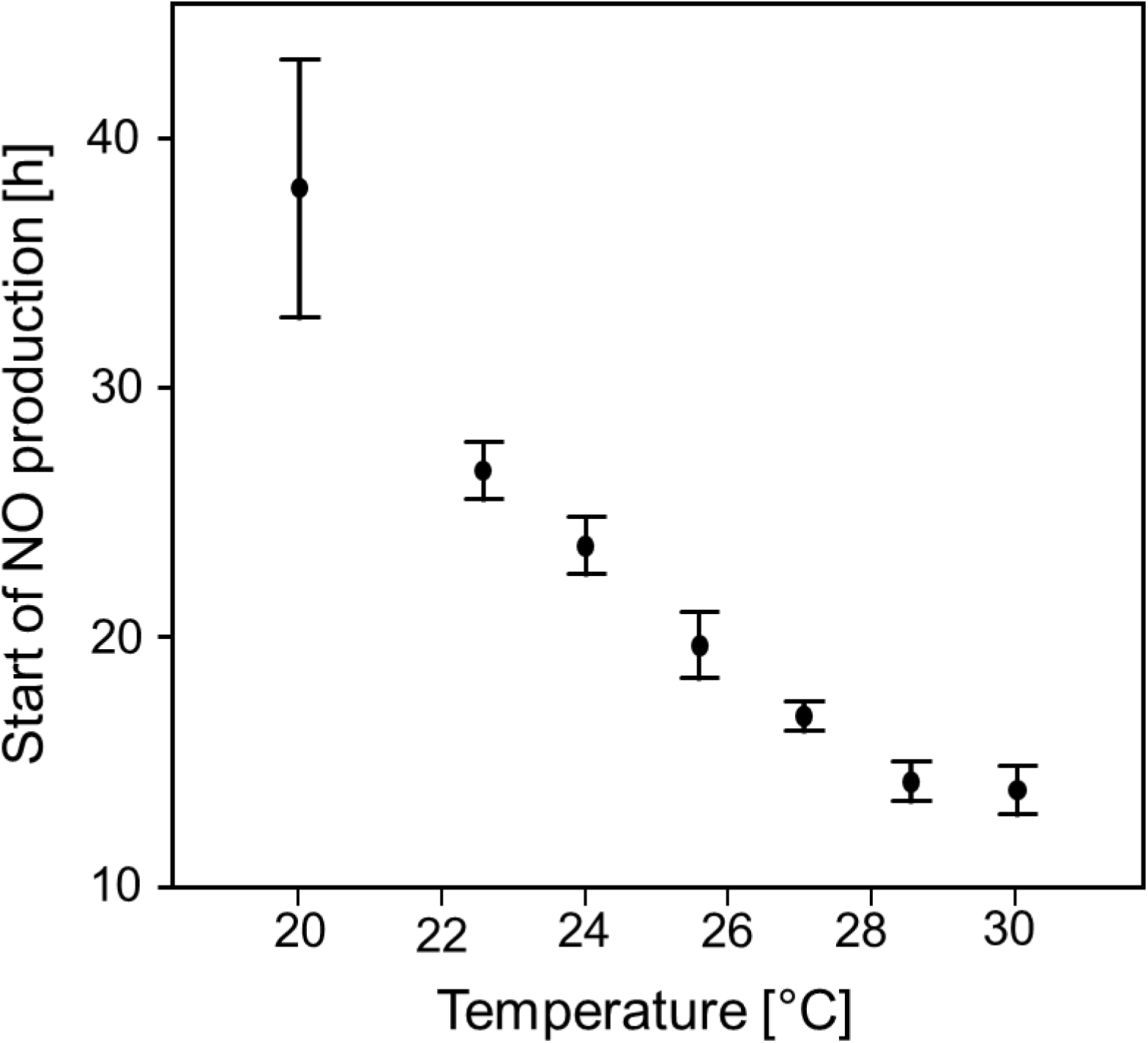
Start of NO˙ emission (h after oviposition) as a function of temperature. Beewolf eggs were kept at different temperatures and the onset of NO˙ release was assessed using the color change of an iodide starch solution as monitored by a digital camera at 30min intervals. Symbols are means ± SD (Quadratic regression: R^2^ = 0.98, N = 33, p < 0.001; Q_10_ = 2.74). Source data file: Fig S2 Source data - start of NO emission.xlsx

**Figure S3.**
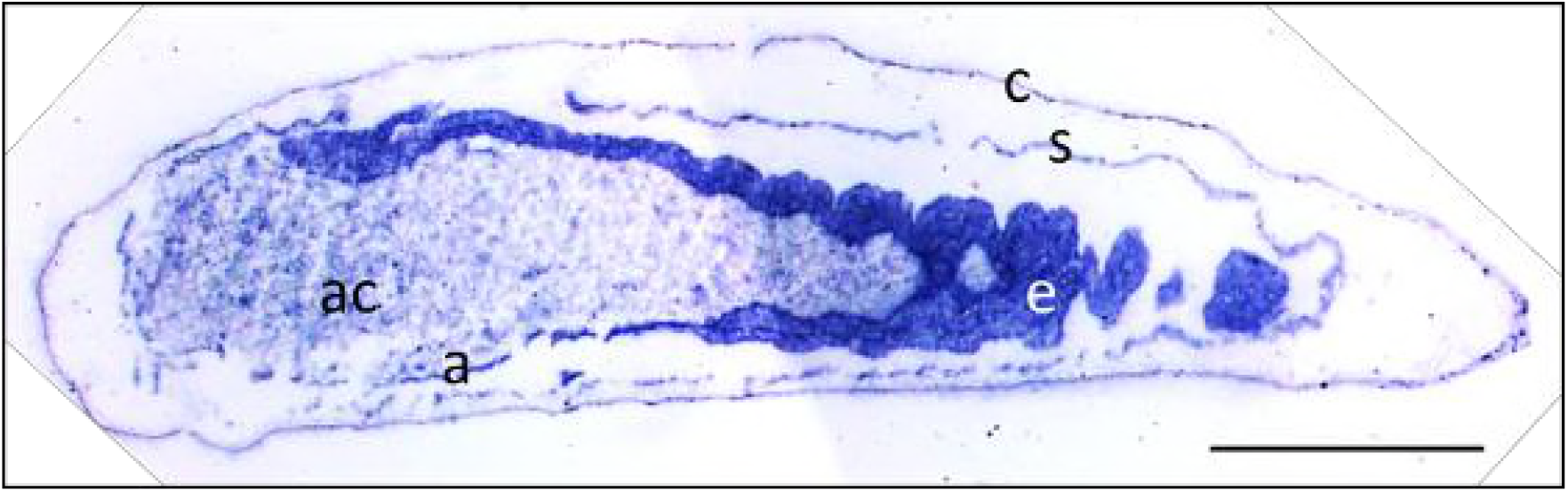
Photomicrograph of a longitudinal section of a beewolf egg showing fixation insensitive NADPH-diaphorase activity. Strong blue staining in the embryonic tissue indicates the presence of reduced nitroblue tetrazolium demonstrating NOS activity (c=cuticle, s=serosa, e= embryo, a=amnion, ac= amnion cavity, scale bar = 1 mm, image composed from two separate photos of the left and right parts of the egg.).

**Figure S4.**
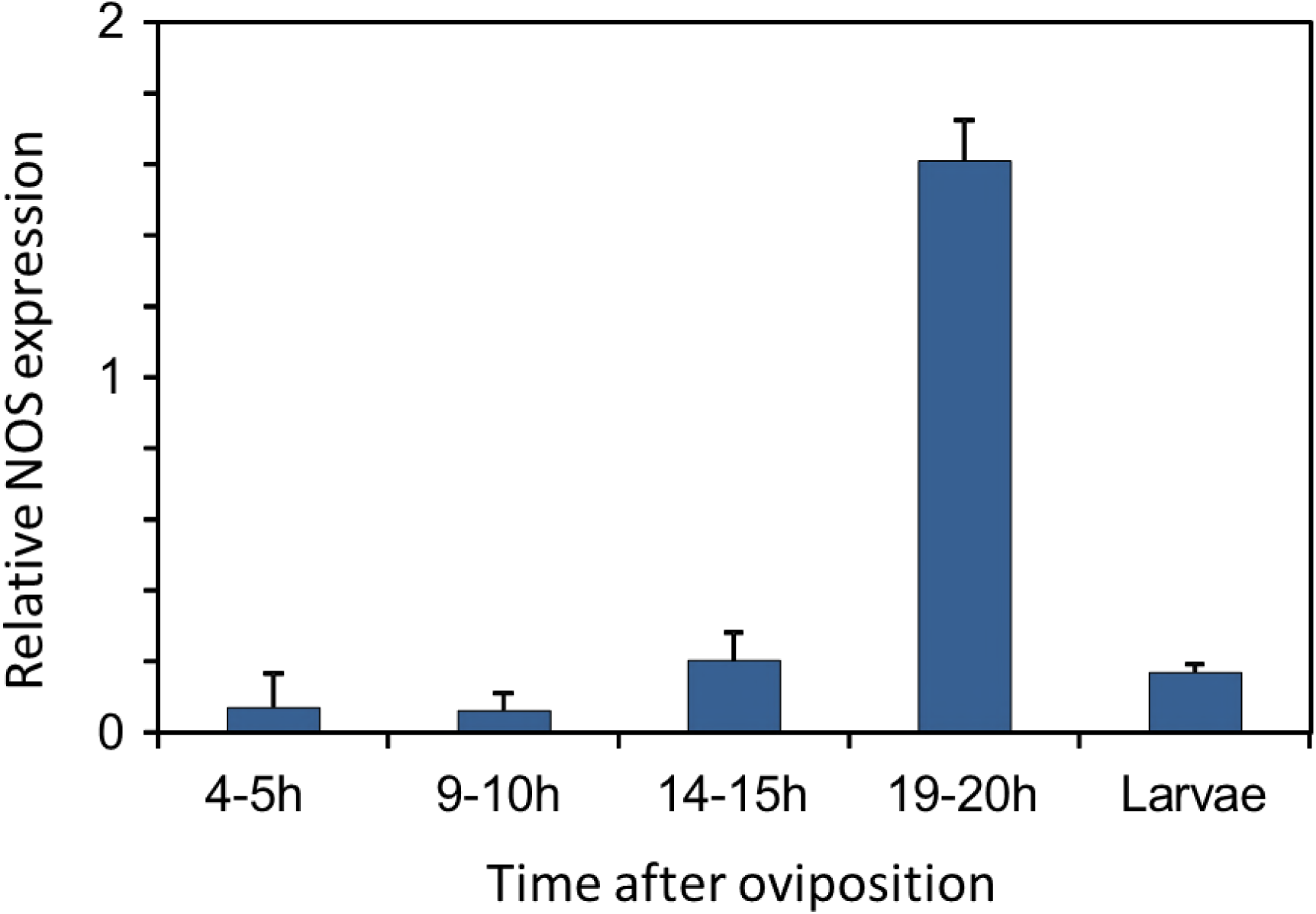
Gene expression of NOS relative to ß-actin in beewolf eggs at different times after oviposition and in freshly hatched larvae. Two trials were conducted, one with 19 and one with 24 eggs per time point. Mean ratios of NOS-mRNA to ß-Actin-mRNA are shown (with standard deviations), as determined by Q-RT-PCR. Source data file: Fig S4 Source data - NOS gene expression.xlsx

**Figure S5.**
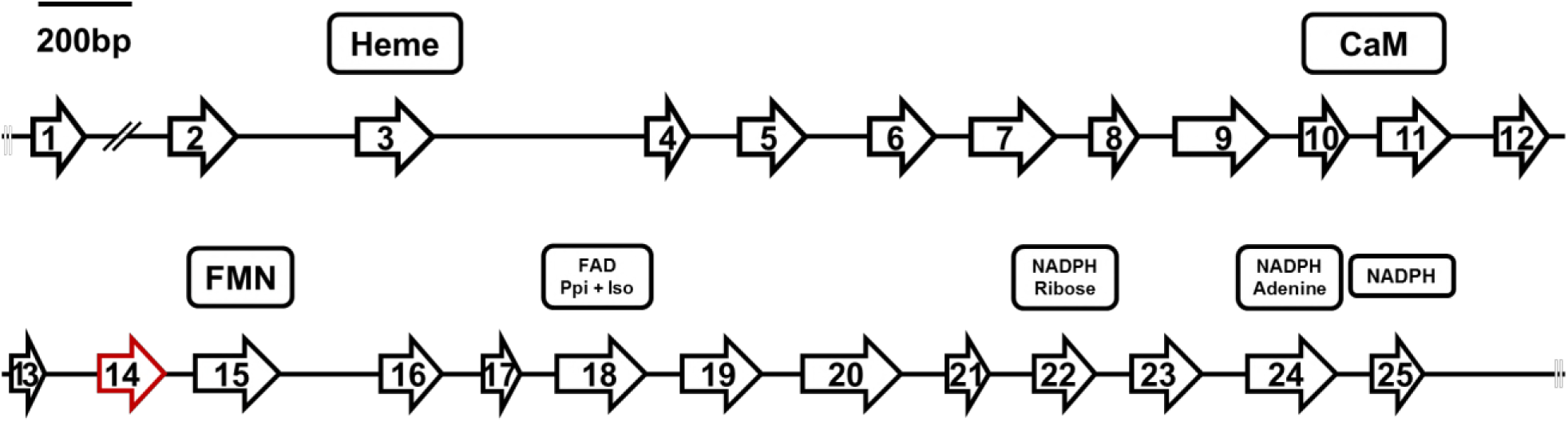
Structure of the *Pt-NOS* gene indicating position and length of exons. Exon 14 (red) is missing in the NOS mRNA in beewolf eggs compared to adults. Presumed cofactor-binding domains as deduced from homologous sequences of the *NOS* of *Anopheles stephensi* [68, 69] are indicated for heme, calmodulin (CaM), FMN, FAD pyrophosphate (FAD PPi) and FAD isoalloxazine (FAD Iso), NADPH ribose, NADPH adenine, and NADPH.

**Figure S6.**
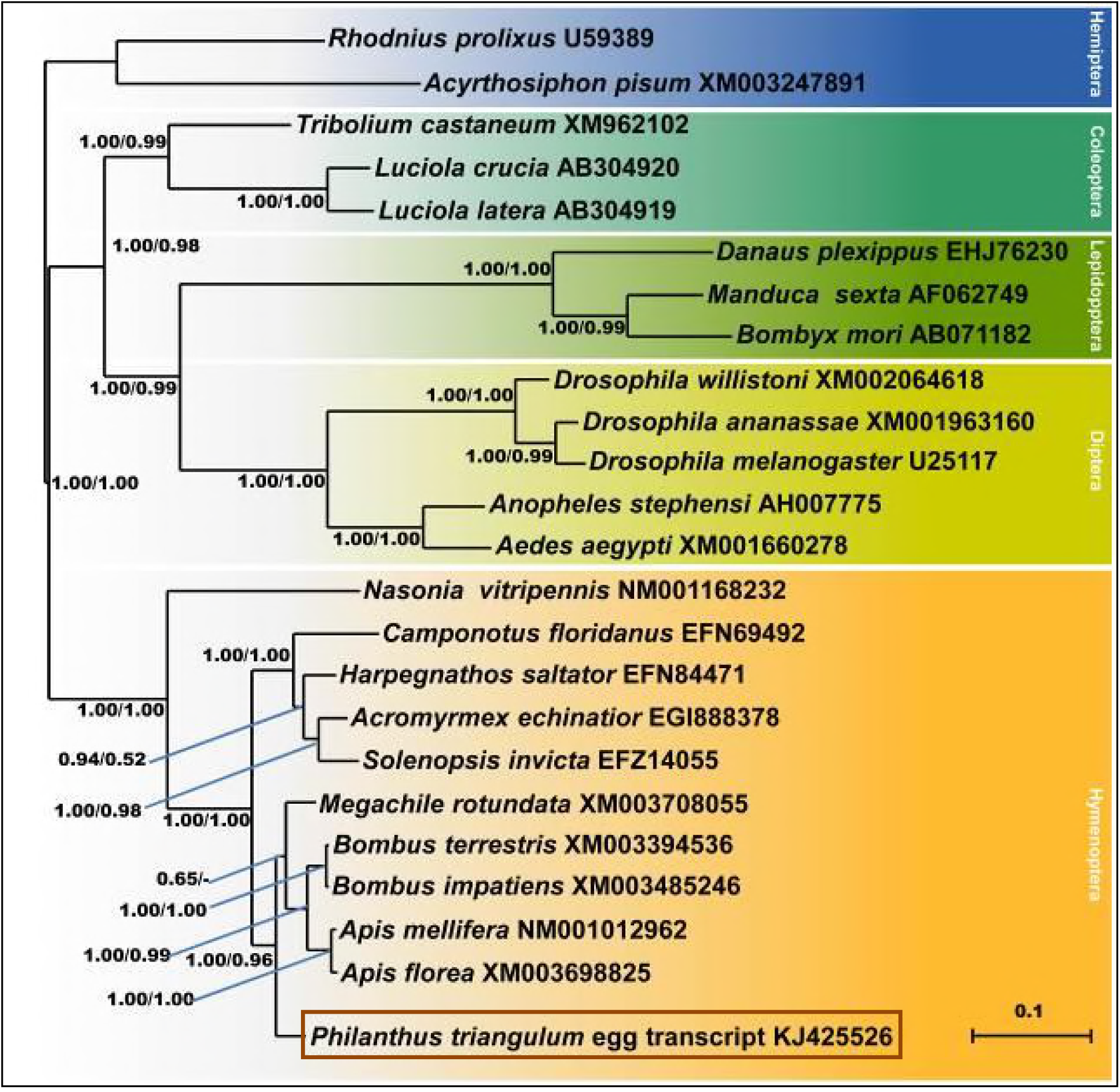
Consensus tree obtained from Bayesian analysis of NOS amino acid sequences from five orders of insects (distinguished by different colors), including the NOS sequences of *P. triangulum* eggs (lowermost entry). Values at the nodes represent Bayesian posterior probabilities and local support values (FastTree analysis), respectively. Scale bar represents 0.1 changes per site.

